# NG2^+^/Nestin^+^ mesenchymal stem cells dictate DTC dormancy in the bone marrow through TGFβ2

**DOI:** 10.1101/2020.10.22.349514

**Authors:** Ana Rita Nobre, Emma Risson, Deepak K. Singh, Julie Di Martino, Julie F. Cheung, Jiapeng Wang, John Johnson, Hege G. Russnes, Jose Javier Bravo-Cordero, Alexander Birbrair, Bjorn Naume, Mohamad Azhar, Paul S. Frenette, Julio A. Aguirre-Ghiso

## Abstract

In the bone marrow (BM) microenvironment, NG2^+^/Nestin^+^ mesenchymal stem cells (MSCs) promote hematopoietic stem cell (HSC) quiescence^1,2^. Importantly, the BM can also harbour disseminated tumour cells (DTCs) from multiple cancers, which, like HSCs, can remain dormant^3^. The BM signals are so growth-restrictive that dormant BM DTCs can persist for years to decades only to awaken and fuel lethal metastasis^3–10^. The mechanisms and niche components regulating DTC dormancy remain largely unknown. Here, we reveal that periarteriolar BM-resident NG2^+^/Nestin^+^ MSCs can instruct breast cancer (BC) DTCs to enter dormancy. NG2^+^/Nestin^+^ MSCs produce TGFβ2 and BMP7 and activate a quiescence pathway dependent on TGFBRIII and BMPRII, which *via* p38-kinase result in p27-CDK inhibitor induction. Importantly, genetic depletion of the NG2^+^/Nestin^+^ MSCs or conditional knock-out of TGFβ2 in the NG2^+^/Nestin^+^ MSCs led to awakening and bone metastatic expansion of otherwise dormant p27^+^/Ki67^−^ DTCs. Our results provide a direct proof that HSC dormancy niches control BC DTC dormancy. Given that aged NG2^+^/Nestin^+^ MSCs can lose homeostatic control of HSC dormancy, our results suggest that aging or extrinsic factors that affect the NG2^+^/Nestin^+^ MSC niche may result in a break from dormancy and BC bone relapse.

---

Metastases, which are derived from disseminated tumour cells (DTCs), are the major source of cancer-related deaths from solid cancers^11^. Ample evidence supports that post-extravasation DTCs can remain in a dormant state from prolonged periods dictating the timing of metastasis initiation^3^. Since years to decades can lapse before dormant DTCs emerge as overt lesions, we postulate that targeting their biology is the shortest path to change patient outcomes by curtailing DTC conversion into metastasis. However, to achieve this goal we must understand the cancer cell intrinsic and micro-environmental mechanisms that control DTC dormancy and reactivation.

The bone marrow (BM) is a common site where dormant DTCs are found and where metastasis can develop in various cancers after prolonged periods of clinical “remission”^3–10^. In trying to understand how the BM microenvironment might control DTC dormancy, we^9,12^ and others^3,5,13,14^ found that in both humans and mice, this microenvironment is a highly restrictive site for metastasis initiation. This is due to the presence of several cues, such as TGFβ2^12^, BMP7^15,16^, GAS6^17–19^ and LIF^14,20^, which induce DTC dormancy in different cancer types. Studies in mostly 2D or 3D *in vitro* models, proposed that mesenchymal stem cells (MSCs)^21^, vascular endothelial cells^13,22^ and/or osteoblasts^18,23,24^ may be the source of dormancy cues for different cancers. However, the function of such niche cells *in vivo* has not been formally tested.

There is a long-standing hypothesis that the niches that control hematopoietic stem cell (HSC) dormancy may instruct DTCs to become dormant^25^. A prior study in prostate cancer has drawn a connection between the HSC niche and dormancy of DTCs^26^. While informative, these studies did not functionally dissect in depth and *in vivo* the role of key cell types that regulate HSC dormancy in regulating DTC quiescence. Further, the niche influence has been inferred from the analysis of cancer cells recovered from the BM or from indirect competition assays. Thus, there is still a critical need to understand the cellular components and how BM niches enforce DTC dormancy.

Dormancy of HSCs in the BM is a robust and long-lived process that, if unperturbed, results in HSCs dividing and self-renewing only 4 to 5 times in the lifetime of a mouse^27^. Such powerful mechanism of quiescence and self-renewal cycles, might explain how DTCs, if responsive to niche signals, could persist for decades in the BM of breast cancer (BC) patients. The HSC microenvironment in the bone marrow is a complex multicellular network promoting HSC dormancy, self-renewal and differentiation into lineage-committed progenitors^2^. Previous studies have revealed that peri-arteriolar stromal cells, enriched in MSC activity, innervated by the sympathetic nervous system, and expressing the neural markers NG2 and Nestin (mesenchymal stem and progenitor cells, hereafter referred to as NG2^+^/Nestin^+^ MSCs for simplicity), are critical for the control of HSC quiescence and hematopoiesis^1^. Importantly, aging-induced alterations causing replicative stress damage or sympathetic neuropathy from infiltrating leukaemia cells, for example, eliminates the control of HSC dormancy by NG2^+^/Nestin^+^ MSCs and can fuel malignancy^28,29^. We and others also discovered that TGFβ2 and BMP7 in the BM milieu are key inducers of DTC dormancy in various epithelial cancers^1,12^. However, the source of these cytokines has remained elusive. Here, we show that NG2^+^/Nestin^+^ MSCs are a source of TGFβ2 and BMP7 and that both the niche MSCs and the cytokines are required to maintain the dormancy of breast and head and neck squamous cell cancer cells (HNSCC) *via* TGFβRIII and BMPRII respectively, p38 and p27 signalling. Furthermore, depletion of the NG2^+^/Nestin^+^ MSCs or knockout of TGFβ2 specifically from NG2^+^/Nestin^+^ MSC compartment led to metastatic outgrowth in the BM without any evidence of inflammation. *In vitro* 3D organoid assays also revealed that “revitalized” but not “aged” NG2^+^/Nestin^+^ MSCs can reprogram malignant cells into a dormancy-like phenotype. Lastly, detectable BMP7 and TGFβ2 levels were observed at higher frequency in the BM of estrogen receptor positive (ER+) BC patients without systemic recurrence during follow up compared to patients with systemic recurrence and BMP7 presence was associated with a longer time to metastatic recurrence after therapy. Our results, pinpoint a functional link between the niches that control adult hematopoietic stem cell quiescence and DTC dormancy.

## Results

### Depletion of NG2^+^/Nestin^+^ MSCs from the BM niche reactivates dormant E0771 DTCs

The BM is a highly restrictive microenvironment for metastasis development but a frequent harbour for dormant DTCs in both humans and mice^3,5,9,12–14^. As observed in humans and prior models of BM DTC detection, in both spontaneous (MMTV-HER2, ^30^ and **Fig1a**) and experimental metastatic BC mouse models (intra-cardiac injected E0771-GFP (**Fig1b**) and MMTV-PyMT-CFP cells) we could only find few DTCs in the BM of femurs, sternum and calvaria bones (150-400 DTCs per million BM cells). In wild type animals these DTCs were non-metastatic or developed lesions at low frequency in the bones (~5-16%, **Fig1e** **and** **4d**). In contrast, these mice showed 100% incidence of metastasis to the lung (data not shown), which unlike the metastasis-restrictive BM niche, is frequently a permissive site for metastasis^12^.

**Figure 1.**
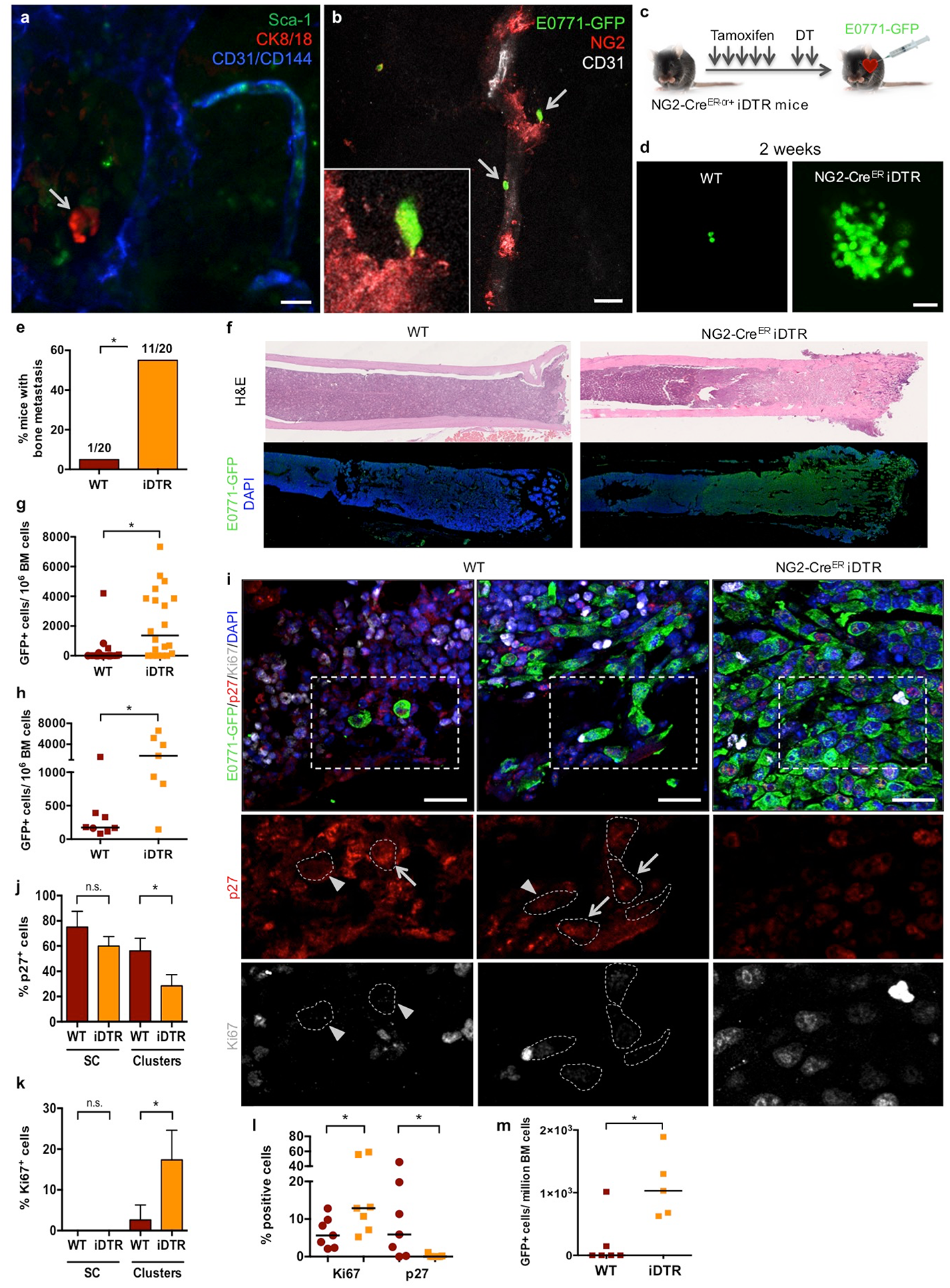
Depletion of NG2^+^/Nestin^+^ MSCs awakens dormant DTCs in the BM. **a-b.** Whole bone imaging of MMTV-Neu (GEM model, CK8/18+ cancer cells) and E0771-GFP (intra-cardiac injected) breast cancer cells. Scale bars: 10um (a), 20um (b). **c-g.** Effect of NG2^+^/Nestin^+^ MSC depletion prior to BM seeding by DTCs (3 independent experiments, n=40, graphs with all mice values and median, 2-tailed Mann–Whitney tests, *p<0.05)**. c.**7-week old NG2-Cre^ER^-iDTR mice (NG2-Cre^ER− or +^) were daily i.p. injected with tamoxifen for 5 days. After 1 rest day, mice were i.p. treated with diphtheria toxin (DT) for 2 days, 24 hours later 2×10^5^ E0771-GFP cancer cells were intra-cardiac injected and 2 weeks later mice were euthanized and the organs collected. **d.** Representative images of E0771-GFP^+^ DTC clusters in BM flushes from WT and NG2^+^/Nestin^+^ MSC depleted mice. Scale bar 50um. **e.** Incidence of bone metastasis (>1000 GFP^+^ DTCs/10^6^ BM cells) 2 weeks after cancer cell injections (Fisher’s exact test, *p<0.05). **f.** H&E and GFP staining of wild type (WT) and NG2-Cre^ER^-iDTR bones. Wild type mice present normal-like bones; while the majority of NG2-Cre^ER^-iDTR mice develop bone metastasis. **g.** Number of E0771-GFP cancer cells per million of BM cells after BM flushing, counted manually. **h.** Number of E0771-GFP cancer cells per million of BM cells after BM flushing, counted by FACS. (n=15, median, 2-tailed Mann–Whitney tests, *p<0.05)**. i.** Representative images of p27 and Ki67 in E0771-GFP cells in the bones. Single cells and small clusters are shown in WT mice and a metastasis in NG2-Cre^ER^iDTR. Scale bars 25um; arrows, positive cells; arrowheads, negative cells; dotted lines, GFP^+^ cells border. **j-k.** Percentage of E0771-GFP cancer cells p27^+^ (**j**) and Ki67^+^ (**k**) detected by immunofluorescence **l.** Percentage of E0771-GFP cancer cells Ki67^high^ and p27^+^ (FACS). (n=14, median, 2-tailed Mann–Whitney U-tests, *p<0.05). **m.** Effect of NG2^+^/Nestin^+^ MSC depletion after BM seeding by DTCs (experimental design in SupFig.1n). Number of E0771-GFP cancer cells per million of BM cells after BM flushing, counted manually (n=10, median, 2-tailed Mann–Whitney U-tests, *p<0.05).

Different factors and BM stromal cells were suggested to play a role in inducing and maintaining DTC dormancy^13,18,21–24^. However, *in vivo* studies functionally linking specific BM cells and dormancy-inducing factors to DTC dormancy are still missing. NG2^+^/Nestin^+^ MSCs are critical inducers of HSC quiescence and hematopoiesis^1^. Thus, we set out to test whether, like dormancy of HSCs^1^, the NG2^+^/Nestin^+^ MSCs could also induce dormancy of DTCs. To this end we took advantage of the C57BL/6 NG2-Cre^ER^iDTR mouse model^1,31^. Treatment of mice with 2 mg of tamoxifen (TAM) for 5 days leads to Cre recombinase activation and the expression of the diphtheria toxin receptor (DTR) in periarteriolar NG2^+^ cells, which upon 2-day treatment with 250ng of diphtheria toxin (DT) causes a targeted depletion of NG2^+^ cells^1,31^. This strategy repeatedly led to the depletion of ~50% of NG2^+^/Nestin^+^ MSCs (which overlap with CD45^−^Ter119^−^CD31^−^PDGFRa^+^CD51^+^ MSCs^1,31^; **Suppl Fig1a,b**) in the BM of long bones. Twenty-four hours after DT treatment, we performed intra-cardiac injection of 2×10^5^ E0771-GFP cells per mouse (**Fig1c** and **Suppl Movie1**) and two weeks later, we monitored for the presence and burden of BM DTCs. All mice had E0771-GFP^+^ DTCs in the BM detectable by FACS (**Suppl Fig1c**), corroborating the injection efficiency and the ability of these cells to persist in the BM at a low burden in all injected mice. Only 5% (1 out of 20) of wild-type (WT) mice showed more than 1000 DTCs/million BM cells (manually counted after BM flush). In contrast, 55% (11 out of 20) of NG2-Cre^ER^iDTR mice with a deficiency of NG2^+^/Nestin^+^ MSCs displayed a dramatic build-up of E0771-GFP^+^ colonies in the BM (**Fig1d,e**). Consistently, histological analysis confirmed that indeed the BM of WT animals exhibited normal histology; while NG2-Cre^ER^iDTR mice revealed large areas of BM metastasis, invasion and replacement of the BM by E0771-GFP^+^ cancer cells (**Fig1f**). Further, quantification of the DTC burden (both manual counting, **Fig1g**, and FACS assisted, **Fig1h**) showed a striking increase in number of DTCs in the BM compartment of NG2-Cre^ER^iDTR compared to WT mice. We conclude that a 50% reduction in NG2^+^/Nestin^+^ MSCs can cause a dramatic expansion of otherwise dormant DTCs into bone-damaging metastasis.

Characterization of proliferation and growth arrest markers revealed that in WT mice, 75% of the solitary E0771-GFP DTCs were positive for p27 (quiescence regulator^12^, **Fig1i,j**) and 54% for pATF2 (p38 activated TF^32^, **Suppl Fig1d,e**). In contrast, these same DTCs were 100% negative for Ki67 and pH3 (**Fig1i,k and Suppl Fig1d,f**), supporting a dormant phenotype (increased expression of pATF2 and p27 and decreased Ki67 and pH3) shown by us and others in different cancer models^12,33,34^. In mice depleted of NG2^+^/Nestin^+^ MSCs, E0771-GFP^+^ resumed proliferation as evidenced by a reduction in the percentage of p27^+^ (28%) and pATF2^+^ (2%) cells and increase of Ki67^+^ (17%) and pH3^+^ (16%) in DTC clusters, which are not frequent in WT animals (**Fig1i-k** **and** **Suppl Fig1d-f**) and corroborated an awakening of DTCs and shift to a proliferative state. Similar results were found by FACS staining of Ki67 and p27 (although the frequencies varied, possibly due to differences between IF and FACS methods) where a significant increase in the percentage of Ki67^+^ E0771-GFP^+^ cells and decrease in p27^+^ DTCs was detected in NG2-Cre^ER^iDTR compared with WT mice (**Fig1l**). When using a MMTV-PyMT-CFP cell line we found a similar trend, only 1 out of 5 (20%) control animals showed greater than 10^3^ cancer cells/10^6^ BM cells, while 3 out of 5 (60%) showed reactivation. The burden of cancer cells in the iDTR group animal that reactivated (60%) had a 10-fold increase in burden and mice with high burden numbers showed a correlative increase in the percentage of Ki67^+^ DTCs upon depletion of NG2^+^/Nestin^+^ MSCs (**Suppl Fig1g-j**).

DT and DTR-mediated cell death can induce inflammation when certain cell types are targeted^35–37^. However, we previously reported no difference in the number of leukocytes, expression of CXCL12 and KITL (SCF) and no change in vascular volume in the NG2-Cre^ER^iDTR model^1^. Nevertheless, because changes could be impacting other inflammatory mediators, we performed a multiplex ELISA for pro-inflammatory cytokines in WT and iDTR mice. These results revealed no differences between the control and iDTR mice in the abundance levels of IL1β, GM-CSF, IL2, IL4, IL6, IL10, IL12p70, MCP-1 or TNFα levels (IFNγ was undetectable - **Suppl Fig2a**). We conclude that the dormant DTC reactivation effect of NG2^+^/Nestin^+^ MSCs depletion is not due to an acute inflammatory response but do to the perturbation of the pro-dormancy HSC niche.

To exclude that the increased DTC burden after two weeks was not due to differences in vascular permeability or extravasation efficiency between WT and NG2-Cre^ER^-iDTR mice, a group of animals was analysed 24 hours after intra-cardiac delivery of E0771-GFP cells and dextran-TexasRed injection (**Suppl Fig2b**). No differences in the amount of 70,000 MW dextran-TexasRed extravasation in the BM was observed after NG2^+^/Nestin^+^ MSC depletion as determined using image analysis (**Suppl Fig2c**), reproducing prior results^1^, and suggesting no obvious alteration in vessel permeability. Further, DTC burden was quantified after expansion of E0771-GFP^+^ cells 1 week *in vitro* (due to no or low detection of E0771-GFP^+^ in fresh BM flush). Similar numbers of E0771-GFP^+^ cells were found in WT and NG2-Cre^ER^-iDTR mice (**Suppl Fig2d**) arguing that the differences in final metastatic burden were not due to higher vessel permeability and/or enhanced extravasation of DTCs.

We next tested whether the depletion of NG2^+^/Nestin^+^ MSC affected DTC expansion once cancer cells had seeded and established a foothold within the NG2^+^/Nestin^+^ MSC niche. To this end we used the same NG2-Cre^ER^iDTR mouse model but the treatment with TAM for 5 days was followed by intra-cardiac injection of E0771-GFP cells, which were allowed to lodge for 72 hours in unperturbed niches and only then we did 2 daily treatments with DT (**Suppl Fig2e**). Importantly, depletion of NG2^+^/Nestin^+^ MSCs after DTCs had lodged in the BM niche also stimulated metastatic outbreak; only a third (1 out of 6) of control WT mice displayed significant expansion of E0771-GFP cells in the BM flushes, while all NG2-Cre^ER^iDTR mice (5 out of 5) had a significant increase in frequency of cancer cells per BM cells (**Fig1m**). We conclude that NG2^+^/Nestin^+^ MSCs are directly or indirectly responsible for maintaining pro-dormancy niches in the BM, which not only control HSC quiescence^1^ but also DTC dormancy.

### NG2^+^/Nestin^+^ MSCs show enhanced production of the dormancy inducers TGFβ2 and BMP7

We had shown that TGFβ2 in the BM was an important cue for dormancy induction^12^, while other groups have pointed for example to BMP7 as a key inducer of DTC dormancy^15^. However, the functional source of TGFβ2 or other factors in the BM *in vivo* remained unidentified. We wondered whether NG2^+^/Nestin^+^ MSC may regulate dormancy by producing cues, such as TGFβ2, that induce DTC dormancy. To address this question, we sorted CD31^−^CD45^−^ Nestin-GFP^bright^ (from now on called Nestin-GFP^+^) and CD31^−^CD45^−^ Nestin-GFP^−^ cells (from now on called Nestin-GFP^−^) stromal cells from mice BM. mRNA levels of TGFβ1, BMP2 (involved in dormant DTC reactivation ^34,38^), TGFβ2 and BMP7 (dormancy inducing cues^12,15,34^) were measured. These experiments showed that Nestin-GFP^+^ cells showed higher mRNAs levels of TGFβ2 and BMP7 than Nestin-GFP^−^ cells (**Fig2a**). Further, ELISA measurements of the BM supernatants from NG2-Cre^ER^iDTR mice showed a significant decrease in TGFβ2 and BMP7 levels but not TGFβ1 or TGFβ3 upon NG2^+^/Nestin^+^ MSC depletion compared with WT mice (**Fig2b**), suggesting that pro-dormancy niches established by NG2^+^/Nestin^+^ MSCs, are a source of TGFβ2 and BMP7 in the BM. Imaging of TGFβ2 and BMP7 in relation to the Nestin-GFP cells revealed that BMP7 was more predominant in the growth plate and appeared to accumulate as a secreted factor around the Nestin-GFP^+^ cells, while TGFβ2 was more homogeneously distributed across the BM, but also in areas containing Nestin-GFP^+^ MSCs (**Fig2c**). We conclude that pro-dormancy NG2^+^/Nestin^+^ MSCs are a source of TGFβ2 and BMP7 that can be detected in the BM microenvironment albeit with varying localization in relation to the NG2^+^/Nestin^+^ MSCs.

**Figure 2.**
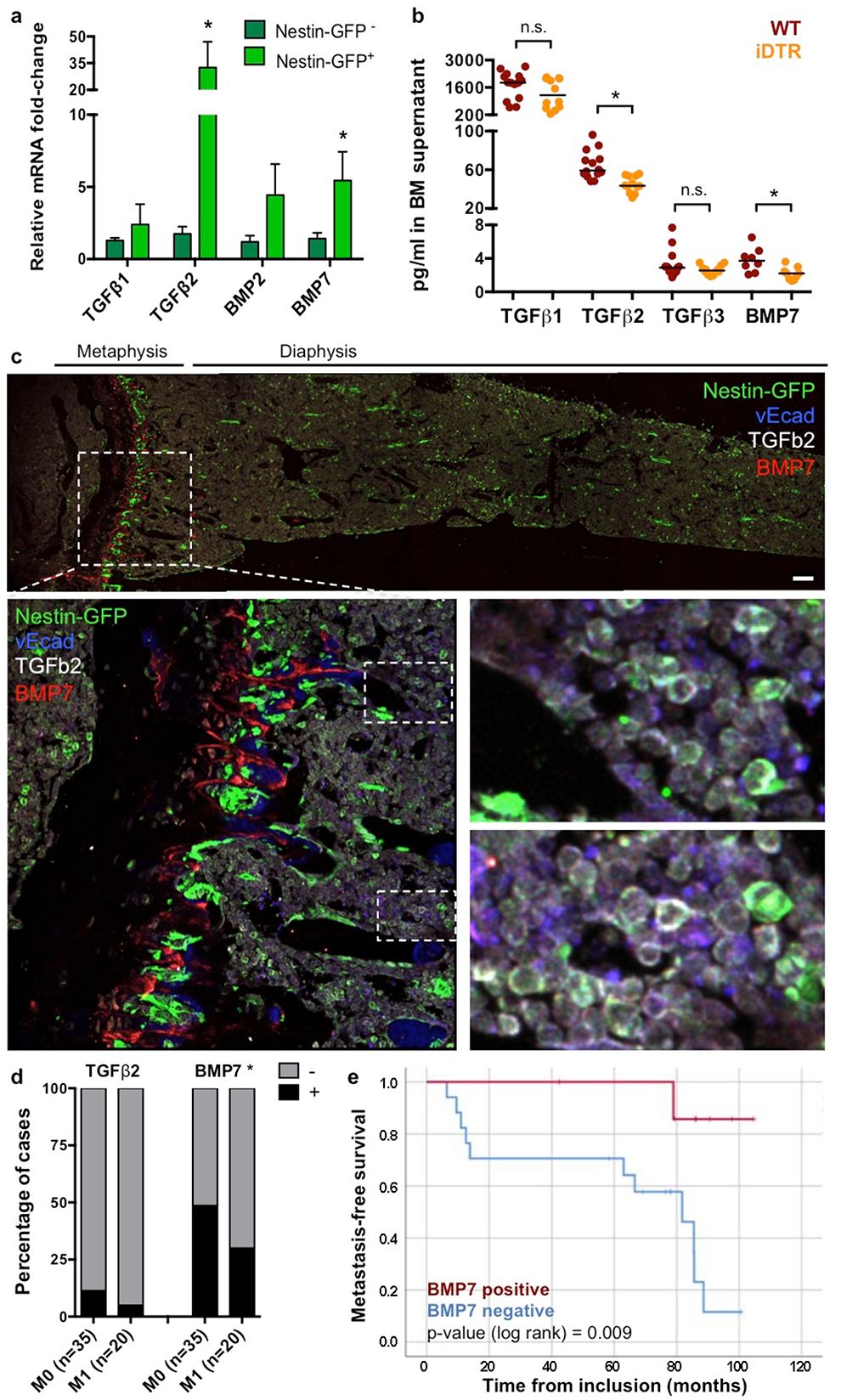
NG2^+^/Nestin^+^ MSCs are a source of pro-dormancy factors TGFβ2 and BMP7 in the BM. **a**. qPCR of sorted CD45^−^CD31^−^Nestin-GFP^−^ and Nestin-GFP^bright^ MSCs from Nestin-GFP mice (4 independent experiments, mean and SEM, 2-tailed Mann–Whitney tests, *p<0.05). **b.** TGFβ1, 2 and 3 and BMP7 levels in BM supernatant of WT and NG2-Cre^ER^-iDTR mice 2 weeks after TAM and DT treatments (2 independent experiments, n=24, median and interquartile range, 2-tailed Mann– Whitney tests, *p<0.05). **c.** Imaging of Nestin-GFP^+^ MSCs in Nestin-GFP mice using IF. Dormancy factors TGFβ2 (white) and BMP7 (red) are expressed near MSCs. Scale bar 100um. Dotted rectangles, high-magnification inserts. **d.** TGFβ2 and BMP7 levels from BM plasma samples from ER^+^ BC patients with (M1) or without (M0) evidence of systemic recurrence (data from SATT clinical study^10^, Fisher’s exact test, *p<0.05). **e.** Metastasis-free survival analysis of the subgroup of patients who did not receive any secondary chemotherapy, excluding treatment-related interpretation bias, separated by detectable BMP7 levels compared to patients with no BMP7.

### TGFβ2 and BMP7 detection in BM supernatant from ER+ BC patients

Given that the lower levels of TGFβ2 and BMP7 upon NG2^+^/Nestin^+^ MSC depletion were associated with reactivation of dormant DTCs leading to metastatic outgrowth (**Fig1**), we tested the hypothesis that the abundance of these two factors in the bone marrow of BC patients might provide some information on their progression. To this end we tested a small subset of BM plasma samples from early BC patients with ER positive disease, the BC subtype most often experiencing tumor dormancy and late recurrences in the bone, among other sites. These samples were part of a clinical study monitoring DTC status after completion of standard adjuvant anthracycline-containing chemotherapy^10^. TGFβ2 and BMP7 levels from BM plasma were detected using multiplex ELISA and revealed that patients without systemic recurrence (M0) during follow up had 2.28- and 1.62-fold higher frequency of detectable TGFβ2 and BMP7, respectively, compared to patients with systemic recurrence (M1) (**Fig2d**). Importantly, metastasis-specific survival analysis of the subgroup of patients who did not receive any secondary chemotherapy, excluding treatment-related interpretation bias, pointed to a markedly improved distant disease-free survival for patients with detectable BMP7 compared to patients with no BMP7 (**Fig2e**) (regardless of the abundance levels). Survival analysis was not possible for TGFβ2 due to the low number of patients showing detectable TGFβ2 levels. While this is a small pilot analysis, these data support further analysis of an association between detectable levels of TGFβ2 and BMP7, which are produced by NG2^+^/Nestin^+^ MSCs, and late recurrences and at least for BMP7, time to metastatic recurrence in ER+ patients after therapy.

### NG2^+^/Nestin^+^ MSCs activate TGFBRIII and BMPRII signalling and a low ERK/p38 signalling ratio in cancer cells

It was not clear if the production of TGFβ2 and BMP7 by NG2^+^/Nestin^+^ MSCs could impact DTC behaviour. To address this question mechanistically, we tested the ability of NG2^+^/Nestin^+^ MSCs to activate key dormancy pathways downstream of TGFβ2 and BMP7 signalling in cancer cells. To this end, we optimized a co-culture system of sorted NG2^+^/Nestin^+^ MSCs and various human and mouse cancer cells. In this assay, Nestin-GFP cells sorted from the BM were co-cultured (1:1 ratio) with cancer cells on top of matrigel at low density. The co-cultures were followed for up to 4 days to ensure that the MSCs retain functionality^39^. We monitored the frequency at which single cancer cells (to mimic solitary DTC biology) remain in a solitary state, or progressed to small and large cancer cell clusters, a measure of proliferative capacity (**Suppl Fig3a** **and** **Fig3a**). The co-cultures revealed that the majority of control (no MSCs added) and also E0771 BC cells co-cultured with Nestin-GFP^−^ cells progressed to large clusters; compared to control cells, Nestin-GFP^−^ cells had some growth inhibitory effect on E0771 cells as we observed ~15% increase in the accumulation of cells in single cell state. However, arrest of cancer cells in a single cell or doublet state was dramatically increased in the presence of Nestin-GFP^+^ MSCs **(Fig3b)**. Similar results were obtained with human head and neck squamous carcinoma T-HEp3 cells **(Suppl Fig3b)** and mouse BC MMTV-PyMT-CFP cells **(Suppl Fig3c)**. Treatment of E0771 cells in these conditions with TGFβ2, BMP7 or BM conditioned media (BM-CM) from WT TGFβ2^+/+^ mice also led to the accumulation of growth arrested single cancer cells **(Suppl Fig3d,h)**. This effect was partially reversed when using BM-CM from mice heterozygous for TGFβ2 (+/−) **(Suppl Fig3h)**. Thus, Nestin-GFP^+^ MSCs, which show enhanced expression of TGFβ2 and BMP7, were able to strongly suppress proliferation of different cancer cells similarly to the purified cues or BM-CM from mice with a full gene complement of TGFβ2. Analysis of the mechanisms revealed that the growth suppression was not due to apoptosis in the co-cultures (measured by c-Cas3 levels), but due to a reduction in p-Rb levels and increase in p-ATF2 (a p38 pathway target) and p27 only observed in the cells co-cultured with Nestin-GFP^+^ MSCs **(Fig3c,f)** or treated with TGFβ2, BMP7 or TGFb2^+/+^ BM-CM **(Suppl Fig3e-k)**. These changes were more evident in solitary E0771 cells, suggesting that in this state cancer cells may be more responsive to dormancy cues. Importantly, TGFβ2 and BMP7 on their own could induce these molecular changes supporting that the Nestin-GFP^+^ MSCs may be activating these pathways through those cues. These results were corroborated using biochemical approaches, which showed that in 2D cultures of E0771 cells TGFβ2 and BMP7 activated SMAD2 and SMAD1/5 phosphorylation respectively and that both converged on the phosphorylation of ATF2 and upregulation of p27 protein levels **(Suppl Fig3l)**, required for dormancy onset^12^. BM-CM from WT TGFβ2 mice led to similar changes in p-SMAD2, p-ATF2 and p27 **(Suppl Fig3l)** as observed in response to TGFβ2 and BMP7; however, BM-CM from mice heterozygous for TGFβ2 (+/−) still activated these pathways suggesting that loss of one TGFβ2 allele cannot fully eliminate the immediate response on the pathway activation within a few hours, but it can reduce the proliferative response over longer periods of treatment **(Suppl Fig3h)**.

**Figure 3.**
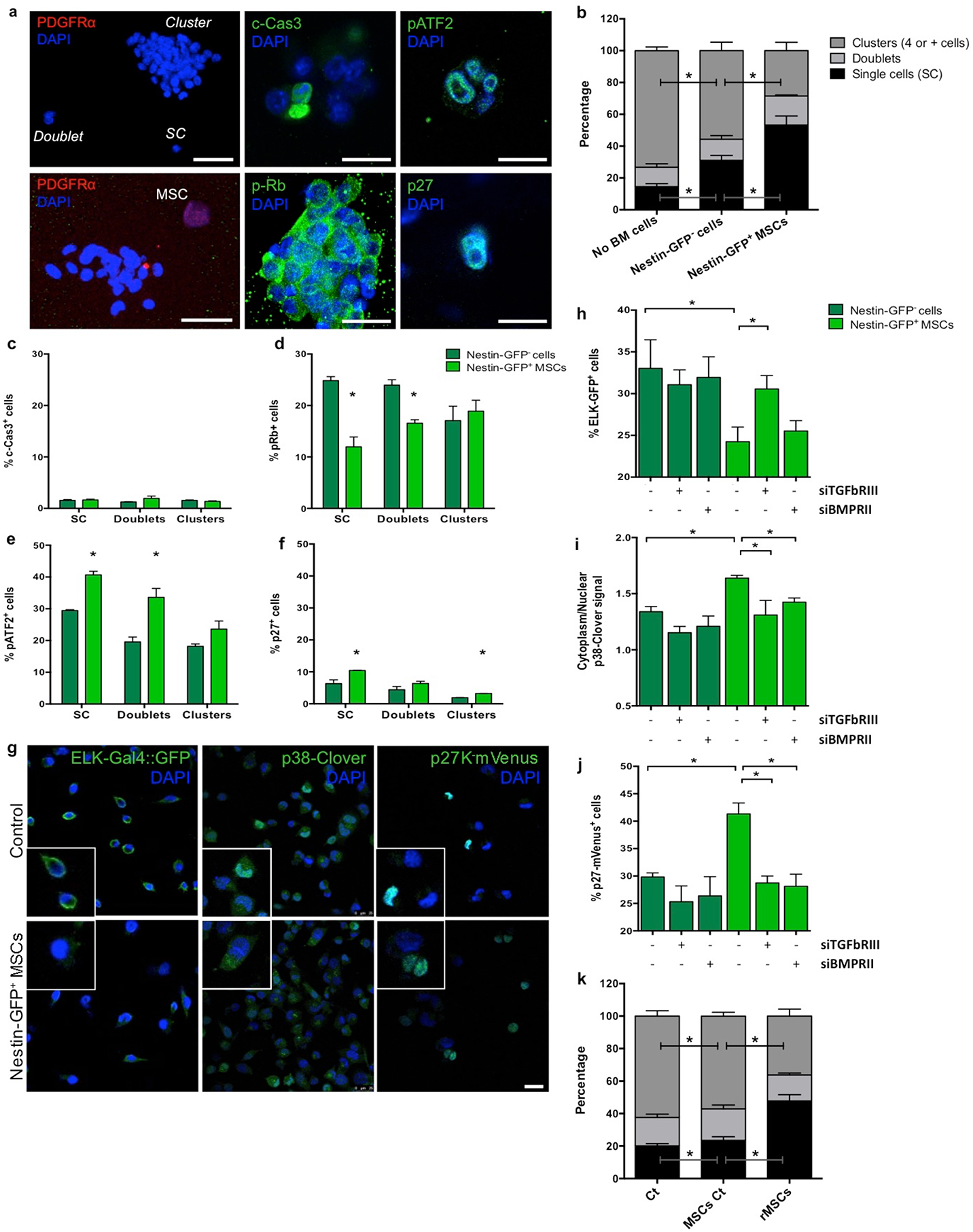
NG2^+^/Nestin^+^ MSCs activate TGFβ2 and BMP7 signalling pathways in cancer cells to inhibit proliferation. **a-f.** Effect of NG2^+^/Nestin^+^ MSCs on tumour cell proliferation. **a.** Representative images of 3D co-cultures of E0771 cells with sorted Nestin-GFP^−^ and Nestin-GFP^+^ MSCs for 4 days. Top left: a single cell, a doublet and cluster of cancer cells. Scale bar 50um; Bottom left: A NG2^+^/Nestin^+^ MSC (PDGFRα^+^, red) near a cancer cell cluster. Scale bar 50um; Centre and right: Representative images of positive cells for cleaved caspase-3 (apoptotic cells), pRb (proliferative cells), p-ATF2 (p38-pathway activation) and p27 (quiescent cells) markers. Scale bars 10um. **b.** Percentage of E0771 cells in a single cell, doublet or cell cluster states after 4 days of co-culture in the indicated conditions. **c-f**. Percentage of E0771 cancer cells positive for c-Cas-3, p-Rb, p-ATF2 and p27 after 4 days of co-culture with Nestin-GFP^−^ cells (dark green) or Nestin-GFP^+^ MSCs (bright green). **g-j.** T-HEp3 cells with different bio-sensors were reversed transfected with siRNA for TGFBRIII and BMPRII followed by co-culture in trans-wells with sorted Nestin-GFP^−^ cells and Nestin-GFP^+^ MSCs. **g.** Representative images of T-HEp3 cells with ELK-Gal4::GFP (GFP^+^ when ERK1/2 pathway is active), p38-Clover (when p38 is active cytoplasmic signal predominates) and p27K^−^ mVenus (mVenus signal indicates cell cycle arrest) bio-sensors. Scale bar 25um. **h-j.** Quantification of the T-HEp3 bio-sensor cell lines after reverse transfection of siRNA for TGFBRIII and BMPRII followed by 24-hour co-culture with Nestin-GFP^−or^ ^+^ cells using trans-wells. **k.** Percentage of E0771 cells in a single cell, doublet or cluster state after co-culture with Control (passaged) or revitalized (r) MSCs for 4 days. All graphs: 3-5 independent experiments, mean and SEM, 2-tailed Mann–Whitney tests, *p<0.05.

Having established that E0771 cells were responsive to TGFβ2 and BMP7 and that Nestin-GFP^+^ MSCs produce these factors, we tested whether the MSC effect was contact dependent and if growth suppression was indeed linked to signalling downstream of these cues. To this end, we used T-HEp3 cells engineered to express an ELK-GAL4::hrGFP biosensor to monitor ERK activity^40^, a p38 shuttle-Clover biosensor that shows clover cytoplasmic over nuclear signal when p38 is active^41^ and a p27K^−^mVenus reporter where protein is stabilized and accumulated in the nucleus upon growth arrest^42^. T-HEp3-biosensor cells were transfected with siRNAs for TGFBRIII and BMPRII and then cultured in the bottom of plates while Nestin-GFP^−^ or Nestin-GFP^+^ cells were plated in upper level of the trans-wells. These experiments showed that Nestin-GFP^−^ cells do not lead to differences in cancer cell ERK, p38 or p27 biosensors activities and that TGFBRIII and BMPRII knockdown (KD) also do not change basal biosensors activities **(Fig3g-j)**. In contrast, Nestin-GFP^+^ MSCs led to significant induction of p38 and p27 activity in cancer cells, even without being in direct contact, and these activations were completely negated by KD of TGFBRIII and BMPRII. Accordingly, Nestin-GFP^+^ MSCs significantly reduced ERK activity in cancer cells, which was also reversed by TGFBRIII, but not significantly by BMPRII knockdown **(Fig3g-j**). Both TGFβ2 and BMP7 activated the p27 and p38 biosensors while inhibiting ELK activity in cancer cells **(Suppl Fig3n-p)**, supporting that they are faithful reporters and that Nestin-GFP^+^ MSCs can directly modulate key dormancy pathways via TGFβ2 and BMP7. In all cases, TGFβ2- or BMP7-dependent effects were reversed by KD of TGFBRIII in the case of TGFβ2 and BMPRII in the case of BMP7 **(Suppl Fig3n-p)**. The above results provide strong evidence supporting that NG2^+^/Nestin^+^ MSCs secrete TGFβ2 and BMP7, to inhibit the mitogenic (ERK1/2) pathway and activate the growth arrest (p38 and p27) pathway.

The ability of long-term cultured NG2^+^/Nestin^+^ MSCs niche cells to maintain HSC self-renewal *ex vivo* is markedly diminished due to loss of the expression of niche factors in cultured MSCs^39^. The loss of these factors is also observed in MSCs passaged 3-5 times in culture^39^ (MSCs Ct); however, revitalized MSCs (rMSCs) engineered to express *Klf7*, *Ostf1*, *Xbp1*, *Irf3* and *Irf7* restore the synthesis of HSC niche factors of BM-derived cultured MSCs^39^. Interestingly, passaged NG2^+^/Nestin^+^ MSCs (MSCs Ct, MSCs kept in culture for 3-5 passages that show loss of function) were not able to suppress growth of E0771 cancer cells. In contrast, rMSCs that have the ability to maintain HSC self-renewal *ex vivo* were able to suppress proliferation of E0771 cells **(Fig3k)**, which correlated with the specific upregulation of TGFβ2 (not TGFβ1) and BMP7 in rMSCs comparted to control MSCs **(Suppl Fig3k)**. These data further support that functional MSCs from the NG2^+^/Nestin^+^ lineage are able to suppress cancer cell growth and that alterations of these cells may impair this dormancy-inducing function.

### Conditional deletion of TGFβ2 in NG2^+^/Nestin^+^ MSCs triggers DTC escape from dormancy in the BM

The above experiments provide strong evidence that NG2^+^/Nestin^+^ MSCs are critical for cancer cell dormancy onset and maintenance in the BM niche (**Fig1**). Additionally, we provide evidence that these MSCs produce significant amounts of TGFβ2 and BMP7 (**Fig2**) and that they activate dormancy signalling pathways in cancer cells (**Fig3**). However, these experiments still do not prove that the dormancy cues derived from NG2^+^/Nestin^+^ MSCs in the niche *in vivo* are required for dormancy. To address this missing link, we focused on TGFβ2 because of our knowledge on the role of this cytokine in dormancy induction and its higher levels in NG2^+^/Nestin^+^ MSCs (**Fig2a** **and**^43^).

We crossed NG2-Cre^ER^ mice with TGFβ2^flox^ mice to generate specific conditional deletion of this cytokine in the NG2 lineage. The NG2-Cre^ER^ driver is mainly restricted to peri-arteriolar Nestin-GFP+ stromal cells as detected using triple-transgenic NG2-Cre^ER^/Nes-GFP/iTdTomato mice; however, it can also be detected in chondrocytes, osteocytes, and rarely in osteoblasts^44,45^. We confirmed decreased levels of TGFβ2 in NG2^+^/Nestin^+^ MSCs (sorted CD45/Ter119/CD31^−^ PDGFRa^+^CD51^+^ MSCs) from the BM of NG2-Cre^ER^TGFβ2 mice tamoxifen induction **(Fig4a-c).** No differences in TGFβ1 levels were found (**Suppl Fig4a**). To test whether disruption of TGFβ2 production in the NG2^+^/Nestin^+^ MSC niche in NG2-Cre^ER^TGFβ2 mice would affect DTC behaviour we injected E0771-GFP cells into the left ventricle of the heart of animals and euthanized them 2 weeks later. All mice had detectable E0771-GFP^+^ DTCs in the BM by FACS (**Suppl Fig4b**), corroborating the injection efficiency. However, only ~16% (4 out of 24) of wild-type (WT) mice shown more than 1000 DTCs/million BM cells (manually counted after BM flush), while 48% (14 out of 29) of NG2-Cre^ER^TGFβ2 (KO) mice displayed a build-up of E0771-GFP^+^ colonies in the BM (**Fig4d**). Consistently, quantification of the DTC burden (both manual counting, **Fig4e**, and by FACS, **Suppl Fig4b**) showed an increase in number of DTCs in the BM compartment of TGFβ2 KO mice compared with WT. Additional analysis revealed that metastatic masses in TGFβ2 KO mice showed a decrease in the percentage of p27^+^ cells (23%) and an increase in the number of Ki67^+^cells (10%) compared to solitary DTCs and small DTC clusters in WT mice (57% p27^+^ / 3% Ki67^+^) **(Fig4f-g)**. A similar difference was found by FACS staining of Ki67, where we found an increase in the percentage of E0771-GFP^+^ Ki67^high^ cells (**Suppl Fig4c**). We conclude that niches containing NG2^+^/Nestin^+^ MSCs and that produce TGFβ2 are required for BC cell dormancy induction and maintenance in the BM compartment.

**Figure 4.**
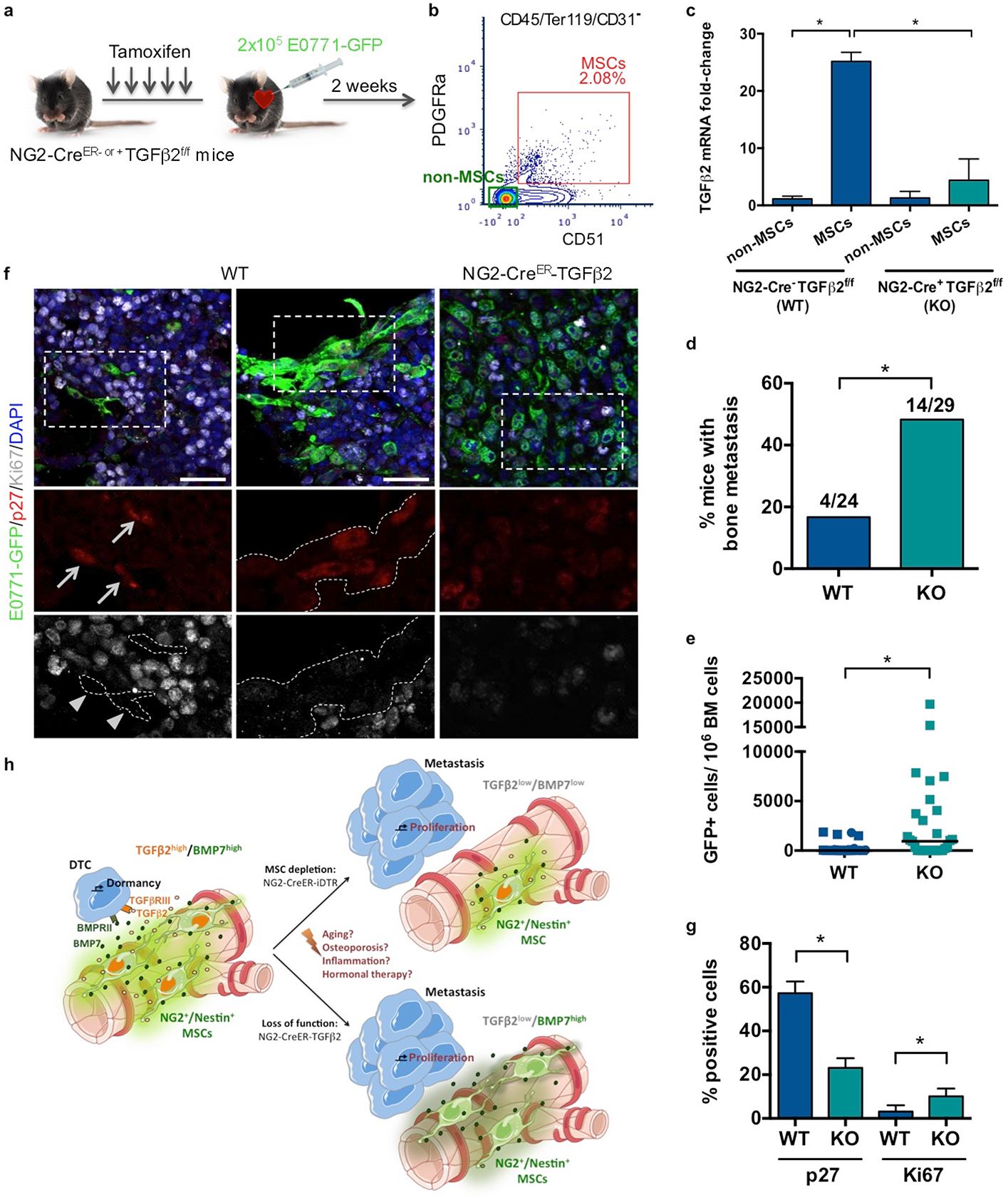
Conditional knock out of TGFβ2 in NG2^+^/Nestin^+^ MSCs awakens dormant DTCs in the BM. **a.** 7-week old NG2-Cre^ER− or +^ TGFβ2^f/f^ mice were i.p. treated daily with tamoxifen (TAM) for 5 days followed by intra-cardiac injection of 2×10^5^ E0771-GFP cells. Mice were euthanized and the organs collected 2 weeks after (4 independent experiments, n=53). **b-c.** Sorting strategy (**b**) and TGFβ2 mRNA levels (**c**) confirming the efficiency of TGFβ2 knockout in NG2^+^/Nestin^+^ MSCs (sorted using CD45^−^Ter119^−^CD31^−^PDGFRa^+^CD51^+^ markers) in NG2-Cre^ER^TGFβ2 mice upon TAM treatments compared with WT mice. **d.** Incidence of bone metastasis (>1000 GFP^+^ DTCs/10^6^ BM cells) 2 weeks after cancer cell injections (Fisher’s exact test, *p<0.05). **e.** Number of E0771-GFP cancer cells per million of BM cells after BM flushing of WT and NG2-Cre^ER^-TGFβ2 mice, counted manually (all mice values and median, 2-tailed Mann–Whitney test, *p<0.05)**. f-g.** Proliferative state of DTCs in the BM of WT and NG2-Cre^ER^-iDTR mice. **f.** Representative images of single cells and small clusters in WT and a metastasis in NG2-Cre^ER^-iDTR. Scale bars 25um; arrows, positive cells; arrowheads, negative cells; dotted lines, GFP+ cells border. **g.** Percentage of E0771-GFP cancer cells p27^+^ (**l**) and Ki67^+^ (**m**) detected by immunofluorescence. **h.** Schematic representation of the current model by which we propose that niches that control HSC dormancy are responsible for inducing dormancy breast cancer DTCs in the BM.

## Discussion

The bone marrow is a powerful pro-cancer dormancy niche for both normal and cancer cells^12–14^. Building on previous studies^13,22^, we show that the BM perivascular niche plays an important role in DTC dormancy. However, here we make a leap in our understanding of this biology by revealing for the first time a specific BM cell type, namely NG2^+^/Nestin^+^ MSCs, that maintains both HSC^1^ and DTC dormancy in this organ. We further advance our understanding by revealing the mechanism behind this biology, where NG2^+^/Nestin^+^ MSCs secrete TGFβ2 and BMP7 to signal through TGFBRIII and BMPRII leading to activation of SMAD, p38 and p27 pathways and cancer dormancy. Importantly, the pro-dormancy function of NG2^+^/Nestin^+^ MSCs is not due to TGFβ2 and/or BMP7 promoting an anti-inflammatory microenvironment. Our work provides novel functional evidence for the long-held notion that HSC dormancy niches support DTC dormancy. Our findings are also deeply linked to the “seed and soil” theory of metastasis^46^. However, our findings reshape this paradigm to show that the “seed and soil” relationship in BM is not for growth of DTCs but rather for the induction and maintenance of dormancy. In fact, our data support that when the “soil” is negatively altered metastasis ensue. Thus, certain organs rather than being a proper “soil” for metastasis may indeed be homeostatic and their normal function as the “soil” is pro-dormancy; when they are altered or damaged they lose their metastasis suppressing function.

Despite our efforts, the low number and density of DTCs in the majority of control mice (150-400 DTCs per million BM cells) in the models we used precluded mapping statistically their location in relation to vascular structures. Thus to gain insight into the relevance of these niches we focused on a functional analysis. We now identify the active role of NG2^+^/Nestin^+^ MSCs in inducing and maintaining DTC dormancy *in vivo*. We went a step further and proved *in vivo* the crucial role of TGFβ2 produced by NG2^+^/Nestin^+^ MSCs. Future studies will test the function of BMP7 in these cells. Our data also suggests that NG2^+^/Nestin^+^ MSCs regulated niches are primarily pro-dormancy rather than only pro-survival, otherwise when the niche was disrupted in the NG2-Cre^ER^-iDTR model, we would see DTC clearance instead of reactivation. However, it is possible that oncogenic signalling in the E0771 cells (K-Ras (activating), MKK4 and p53 (both inactivating mutants)^47^ protects them from absent survival signals that could be altered when we eliminate NG2^+^/Nestin^+^ MSCs. It is interesting that even carrying MKK4 (p38 and JNK upstream activator) and p53-inactivating mutations (and K-Ras active mutant) E0771 efficiently activated p38 signalling and CDK inhibitors long term. This suggests that TGFβ2 and BMP7 signalling may be more reliant on MKK3/6 signalling for p38 activation. It also supports that niche cues can override potent oncogene signalling, further solidifying the notion that microenvironmental mechanisms may be dominant over genetics^48,49^ if cancer cells can still “read” host homeostatic signals.

We also report no major differences in vessel permeability and DTC early seeding after NG2^+^/Nestin^+^ MSCs depletion, supporting that the enhanced DTC growth in NG2^+^/Nestin^+^ MSCs depleted before or after seeding in the BM was due to a dormancy-to-growth switch effect and not a seeding/colonization phenotype. An important next question will be to define how cancer cells with different genetics and from different cancer types might affect the HSC niche to support or block a positive loop feeding dormancy of both HSC and DTCs in an homeostatic normal-like BM environment. Our results may also start to define what genetic alterations may allow cancer cells to escape microenvironmental control by NG2^+^/Nestin^+^ MSCs. For example, genetic alterations that affect TGFBRIII^50–52^ and BMPRII^38,53^ have been linked to increased bone metastasis in breast and prostate cancer, indicating that such alterations may allow DTCs to escape homeostatic control. These genetic changes detected in DTCs may serve as biomarkers to help monitor more closely patients at risk of relapse. To this end we provide promising data that monitoring for example BMP7 presence or absence in ER+ breast cancer patients can inform on the time to metastatic relapse. Patients that presumably had intact or functional NG2^+^/Nestin^+^ MSC and/or other niches and had detectable BMP7 were less prone to develop metastasis.

It is important to highlight the complexity of the BM niches and the fact that different cell types may produce the same or other dormancy cues, which we observed in our co-cultures where Nestin-GFP^−^ cells led to a slight increase in the percentage of cancer cells in a single cell state (Fig3b). Also, the driver we used NG2-CreER is mainly restricted to MSCs, but previous studies in triple-transgenic NG2-cre^ER^/Nes-GFP/iTdTomato mice (same driver) showed that TdTomato expression was observed mainly in peri-arteriolar Nes-GFP+ stromal cells and to some extent in chondrocytes and osteocytes. However, TdTomato rarely marked osteoblasts even after 5 months^44^ (and unpublished data) as previously reported by Zhou *et al.*^45^. Overall, it was in fact rather surprising that a relatively rare subpopulation of MSCs had such a powerful effect on DTC dormancy *in vivo*. This could be due to the fact that DTCs in the BM are not overwhelming the niches due to low abundance and/or that the NG2^+^/Nestin^+^ MSCs coordinate pro-dormancy niches by instructing other cell types to produce dormancy cues. This is possible as other cell types in the BM niche are also key contributors to the pro-dormancy HSC niche. To this end, it was previously described that osteoblasts are also a source of TGFβ2^24,43^, but in our mouse models these cells do not seem to compensate for the loss of TGFβ2 in NG2^+^/Nestin^+^ MSCs or for the 50% loss of NG2^+^/Nestin^+^ MSCs after DT treatment. However, it is possible—but not tested—that osteoblasts are producing TGFβ2 also in response to niche signals orchestrated by NG2^+^/Nestin^+^ MSCs. NG2^+^/Nestin^+^ MSCs may also produce additional factors to TGFβ2 and BMP7, which would explain why the penetrance of the phenotype seems greater when we compare DTC burden in the NG2-Cre^ER^-iDTR model (Fig1) vs. the NG2-Cre^ER^-TGFβ2 (Fig4).

In our models we observe that while the HSC dormancy niches robustly control the dormant DTC phenotype, when we deplete NG2^+^/Nestin^+^ MSCs, growth is alarmingly aggressive. This argues that identifying the key niche cells that maintain dormancy in patients may be important to monitor their function to predict and prevent relapse. With this in mind, our current results and those from past studies argue that it may be more feasible to eradicate dormant DTCs or maintain them in dormancy than to awaken them and try to kill them with conventional therapies; their growth post-awakening is aggressive and may occur in multiple organs^12^.

It is known that aging affects the proper function of HSC niches by downregulation of niche factors by passaged MSCs^39^ and that in aged mice the BM microenvironment shows lower levels of TGFβ2 and BMP7^54^. Our results offer a new opportunity to understand how aging, inflammation and the cancer cells themselves may alter MSCs functionality leading to a disruption of the BM pro-dormancy niche that ultimately leads to dormancy awakening and bone metastasis formation. Finally, a remaining question, but beyond the scope of our study, is whether the function of NG2^+^/Nestin^+^ MSCs as pro-dormancy niche orchestrators is limited to the BM or whether it has similar roles in other metastasis target organs. Overall, we propose that this work shifts our understanding on how homeostatic mechanisms that control adult stem cell quiescence may govern dormancy of disseminated cancer cells and control the timing of metastasis. Our work reshapes the paradigm of metastasis by revealing how the homeostatic BM microenvironment actually serves mainly as a metastasis suppressive “soil” via dormancy induction.

---

**Supplementary Figure 1.**
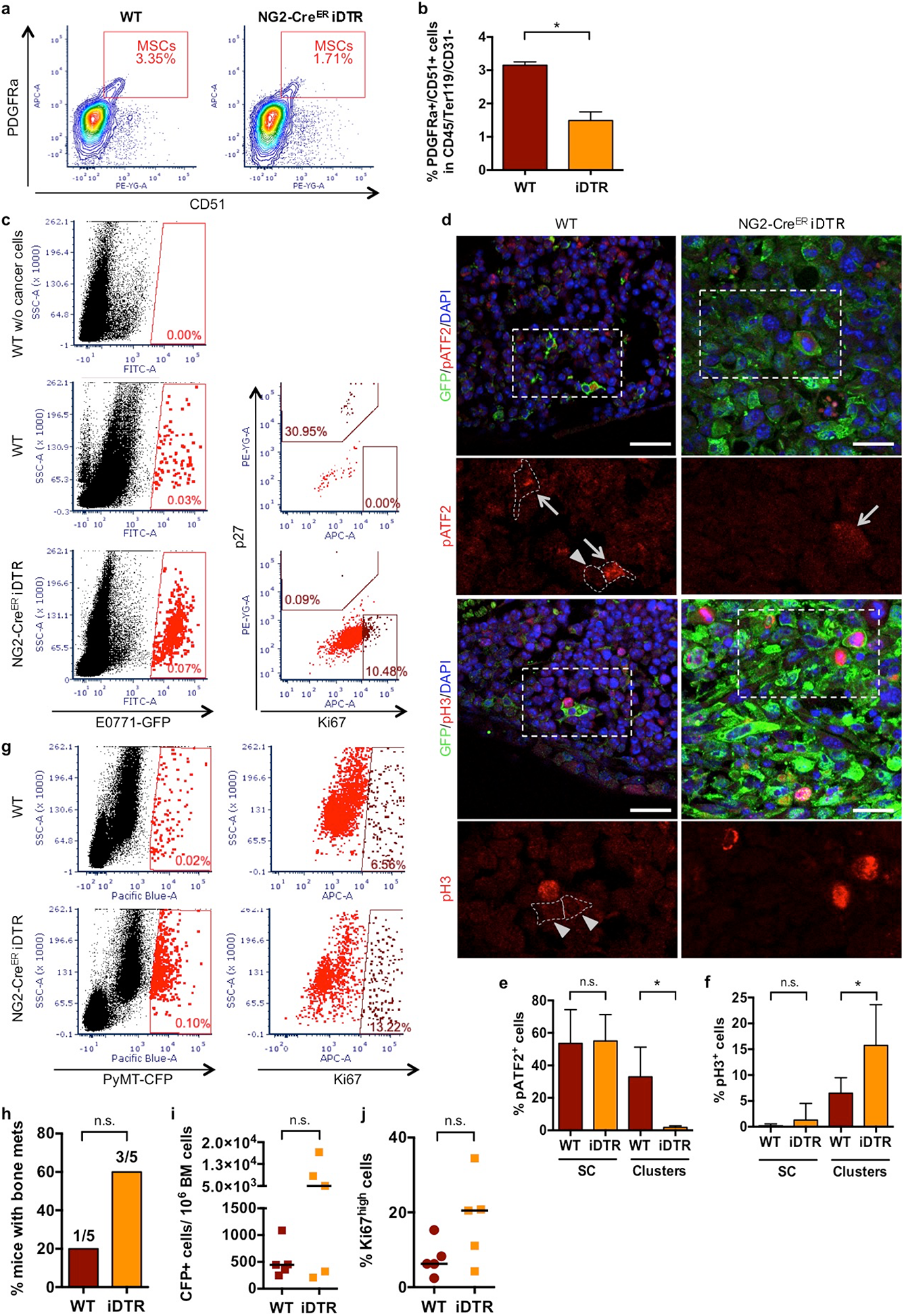
Controls for depletion of NG2^+^/Nestin^+^ MSCs, DTC presence in the BM and their characterization. **a-b.** FACS plots (**a**) and quantifications (**b**) confirming the depletion of NG2^+^/Nestin^+^ MSCs (CD45^−^Ter119^−^CD31^−^PDGFRa^+^CD51^+^) in NG2-Cre^ER^iDTR mice upon TAM and DT treatments compared with WT mice. **c.** Representative plots and gates used in FACS for detection and characterization of E0771-GFP cells in BM flushes. **d-f.** Representative images (**d**, Scale bars 25um; arrows, positive cells; arrowheads, negative cells) and quantification of pATF2^+^ (**e**) and pH3^+^ (**f**) E0771-GFP DTCs. **g-j.** Detection and characterization of MMTV-PyMT-CFP cells in BM flushes by FACS (n=10). **h.** Incidence of bone metastasis (>1000 GFP^+^ DTCs/10^6^ BM cells) 2 weeks after cancer cell injections (Fisher’s exact test, *p<0.05). **i.** Number of MMTV-PyMT-CFP cancer cells per million BM cells. **j.** Percentage of Ki67^high^ E0771-GFP cancer cells. All graphs include all mice values, median, 2-tailed Mann–Whitney U-tests, *p<0.05.

**Supplementary Figure 2.**
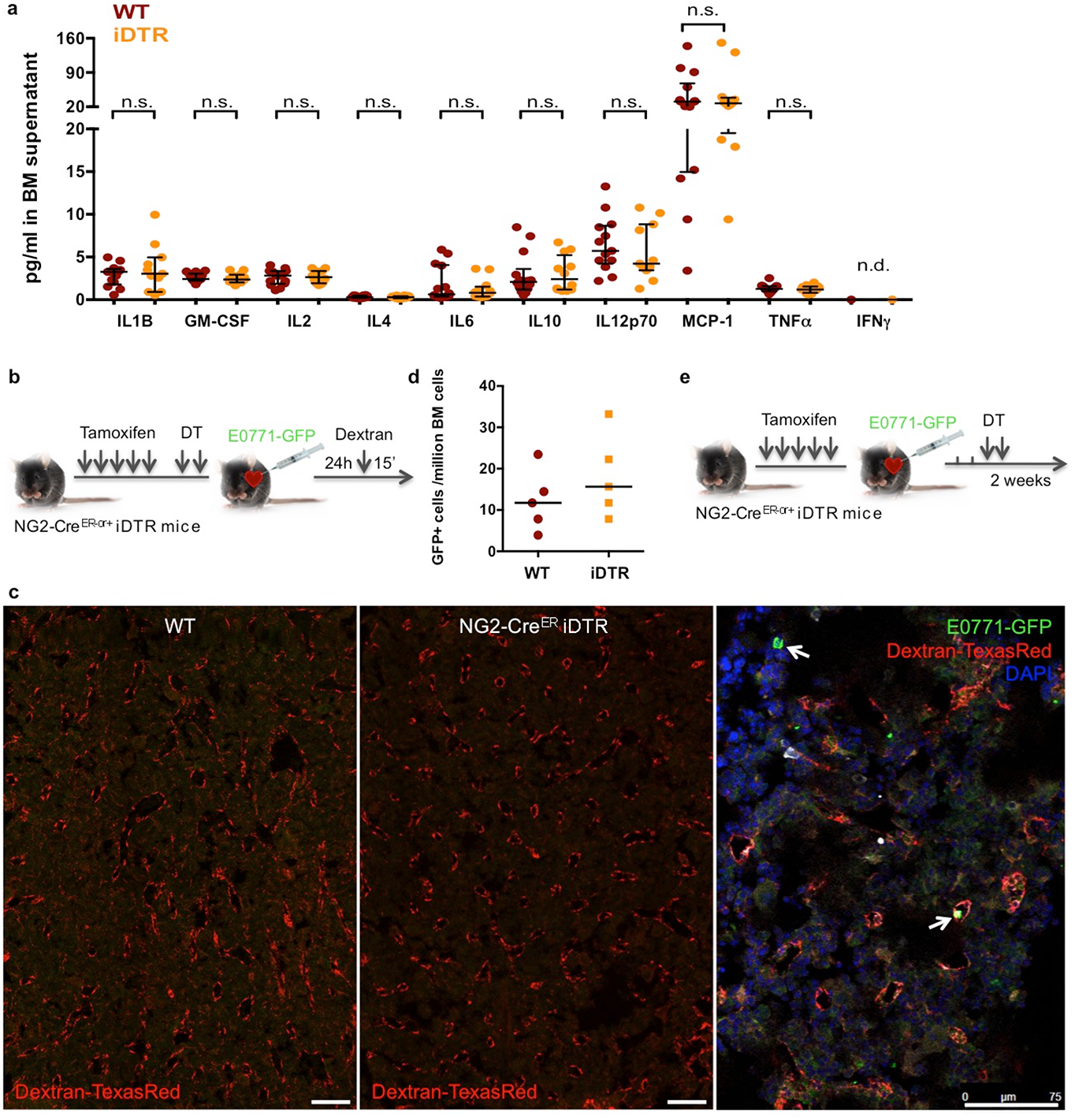
Pro-inflammatory cytolkine status, vessel permeability and BM DTC seeding in NG2^+^/Nestin^+^ MSCs-depleted mice. **a.** Levels pro-inflammatory cytokines in BM supernatant of WT and NG2-Cre^ER^-iDTR mice 2 weeks after TAM and DT treatments (n.d. not detected; 2 independent experiments, n=24, median and interquartile range, 2-tailed Mann–Whitney tests, *p<0.05). **b-d.**7-week old NG2-Cre^ER^-iDTR mice (NG2-Cre^ER− or +^) were daily i.p. injected with tamoxifen for 5 days followed by a rest day and 2 i.p. injections of DT. 24 hours later, 2×10^5^ E0771-GFP cancer cells were delivered via intra-cardiac injection and 24 hours after injected with 70K Dextran-TexasRed, 15 minutes prior euthanasia (n=10). **c.** Representative images of Dextran extravasation in perfused bones. No differences in Dextran extravasation were detected between WT and NG2-Cre^ER^-iDTR mice arguing for a lack of effects on vascular permeability. E0771-GFP cancer cells (arrows) were detected in both WT and NG2-Cre^ER^-iDTR mice. **d.** Number of E0771-GFP cells detected after 1 week of *in vitro* expansion of the BM aspirates collected 24 hours after injection into mice. Similar numbers were detected in WT and NG2-Cre^ER^-iDTR mice supporting no differences in seeding capacity. **e**. 7-week old NG2-Cre^ER^-iDTR mice (NG2-Cre^ER− or +^) were daily i.p. injected with TAM for 5 days, followed by intra-cardiac injection of 2×10^5^ E0771-GFP cancer cells. Cells were allowed to disseminate and extravasate for 72 hours followed by 2 i.p. injections of DT. 2 weeks after cancer cell injection mice were euthanized and the organs collected.

**Supplementary Figure 3.**
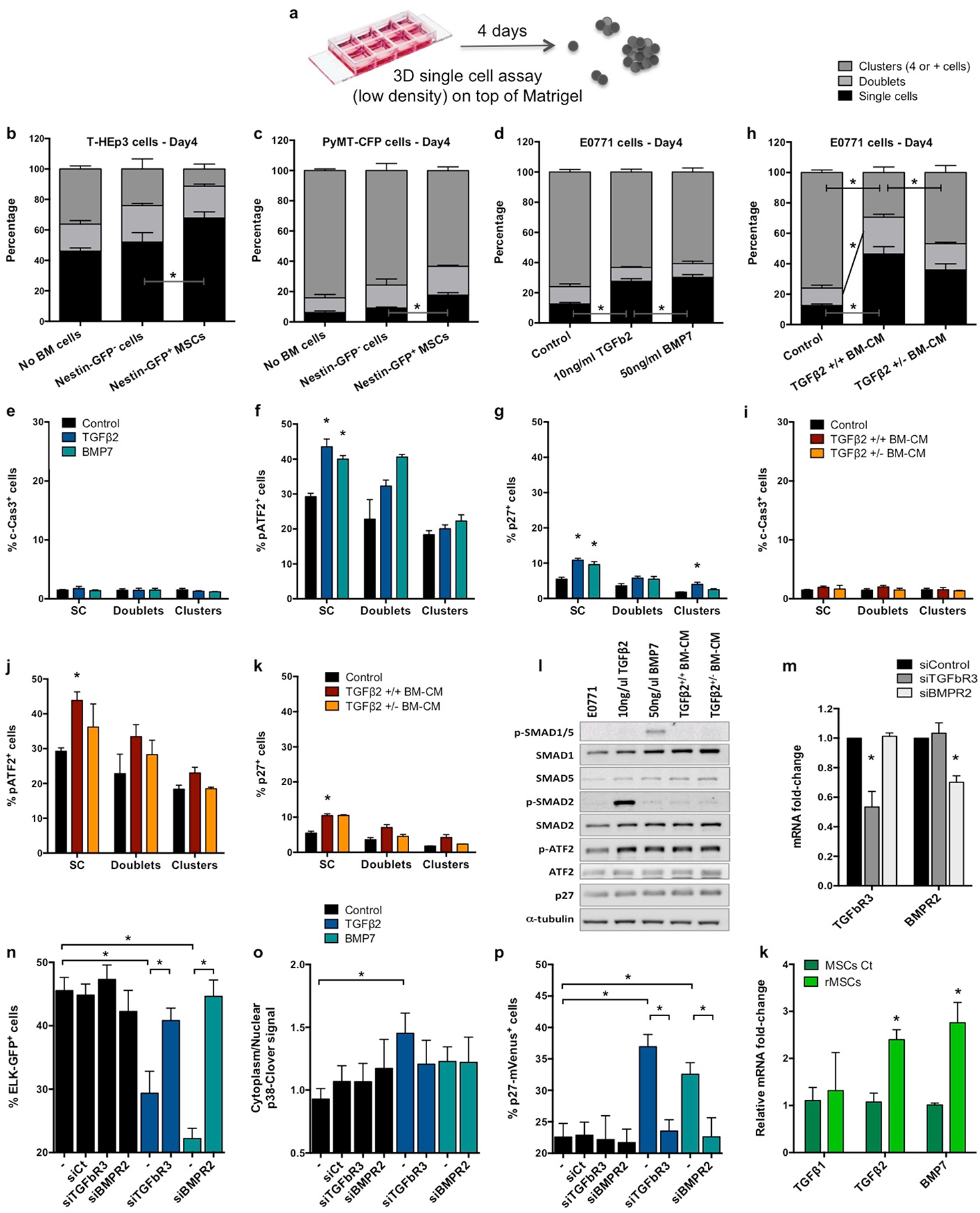
Effect of NG2^+^/Nestin^+^ MSCs on signalling pathways and growth of various cancer cell types in 2D and 3D cultures. **a.** 3D Matrigel assay used to track solitary cell to cluster growth. Single cells were plated on top of Matrigel in low density and the percentage of single cells, doublets and clusters was quantified 4 days after. **b-c.** Co-culture of human HNSCC PDX-derived T-HEp3 (**b**) and mouse BC MMTV-PyMT (**c**) cells with sorted Nestin-GFP^−^ and Nestin-GFP^+^ MSCs for 4 days. **d-k.** E0771 cells were treated every day for 4 days with TGFβ2, BMP7 or bone-marrow conditioned media (BM-CM) of TGFβ2^+/+^ or TGFβ2^+/−^ mice. **d and h.** Percentage of cancer cells in a single cell, doublet or cluster state with the indicated treatments. **e-g and i-k.** Quantifications of c-Cas-3, p-ATF2 and p27 with the indicated treatments. **l.** Western blots for the indicated antigens detected in E0771 cells treated for 24 hours with the TGFβ2, BMP7 and different BM-CM preparations. **m-p.** T-HEp3 cells with ERK, p38 and p27 activity biosensors were reversed transfected with control siRNA or siRNAs for TGFBRIII and BMPRII followed by 24-hour treatments with TGFβ2 and BMP7. **m.** TGFBRIII and BMPRII mRNA levels 48 hours after transfection with the indicated siRNAs. **n-p.** Quantification of the T-HEp3-biosensors activity. **k.** qPCR of TGFβ1, TGFβ2 and BMP7 from Control and rMSCs. All graphs: 3-5 independent experiments, mean and SEM, 2-tailed Mann–Whitney tests, *p<0.05.

**Supplementary Figure 4.**
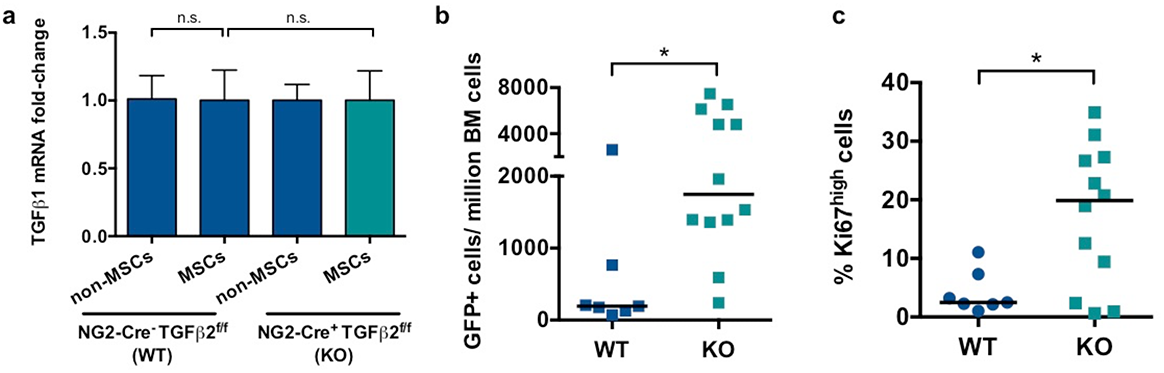
NG2-Cre^ER^TGFβ2 mouse model controls. **a.** TGFβ1 mRNA levels in NG2^+^/Nestin^+^ MSCs (sorted using CD45^−^Ter119^−^CD31^−^PDGFRa^+^CD51^+^ markers) in NG2-Cre^ER^TGFβ2 mice upon TAM treatments compared with WT mice. **b.** Number of E0771-GFP cancer cells per million of BM cells after BM flushing, counted by FACS. **c.** Percentage of Ki67^high^ E0771-GFP cancer cells (FACS). All graphs include all mice values, median and interquartile range and 2-tailed Mann–Whitney U-tests, *p<0.05.

**Supplementary Movie1**– movie of a typical sonogram used to do imaging guided injection of cells in the left ventricle of the heart. This methodology ensures intra-cardiac injection efficiency, minimizing variability of injections on the BM DTC burden analysis.

## Methods

### Animals

Nes–GFP^55^ were bred in and obtained from the Frenette’s laboratory at Albert Einstein Institute. B6.Cg-Tg(Cspg4-cre/Esr1*)BAkik/J (NG2-Cre^ER^ - Jackson Laboratory, stock 008538), C57BL/6-Gt(ROSA)26 Sortm1(HBEGF) Awai/J (iDTR – Jackson Laboratory, stock 007900) and Tgfβ2^flox^ (^56^, gift from Mohamad Azhar’s lab) were maintained on the C57BL/6J background and bred and crossed in our facilities. All NG2-Cre^ER^iDTR and NG2-Cre^ER^TGFβ2 mice were genotyped using the primers in **Supplementary Table 1**. 7 to 8-week-old female mice were used. No randomization or blinding was used to allocate experimental groups. All experimental procedures were approved by the Institutional Animal Care and Use Committee (IACUC) of Icahn School of Medicine at Mount Sinai.

### Induction of NG2-Cre^ER^-mediated recombination and iDTR-mediated cell depletion

7 to 8-week old female mice were injected intraperitoneally with 2mg of tamoxifen (Sigma, T5648) dissolved in corn oil daily for 5 days to induce the Cre^ER^-mediated recombination. In case of NG2-Cre^ER^-iDTR, mice were also injected intraperitoneally with 250ng of diphtheria toxin (Millipore, 322326) dissolved in serum chloride daily for 2 days for iDTR-mediated cell depletion.

### Metastasis assays

2×10^5^ E0771-GFP or MMTV-PyMT-CFP cells were intra-cardiac injected with echo-guidance using micro ultrasound Vevo2100, Transducer MS-250, 21MHz and Vevo LAB 3.1.1 software (FUJIFILM VisualSonics Inc.) **(Supplementary Movie 1)**. 24 hours or 2 weeks later, mice were euthanized and organs were collected and processed. In a group of mice, 70,000 MW Lysine fixable Dextran Texas Red (Invitrogen) was retro-orbital injected 15 minutes prior to euthanasia and mice were perfused with PBS. Bone marrow (BM) cells were flushed from femurs and tibias, red-blood-cell lysis buffer (Lonza) was used for 2 minutes followed by quantification of the GFP+ cells and normalization to the total number of BM cells.

### Flow cytometry and cell sorting

Bone marrow cells were flushed, incubated in red-blood cell lysis buffer (Lonza) for 2 minutes and remaining cells were permeabilized with 0.05% Triton (when using intracellular antibodies) and stained using antibodies and conditions in **Supplementary Table 2**. All experiments were performed using BD FACSAria II equipped with FACS Diva software (BD Biosciences). Dead cells and debris were excluded by FCS, SSC and DAPI (4’,6-diamino-2-phenylindole) (Fisher Scientific) staining profiles. Data were analysed with FACS Diva (BD Biosciences) or FCS Express Cytometry 7 (De Novo) softwares.

### Histopathology

After dissection, bones were fixed in 4% paraformaldehyde (PFA, Thermoscientific) for 24 hours. Bones were then decalcified in 14% EDTA, renewed every other day for 7-10 days at 4°C with agitation. Bones were processed, embedded in paraffin and sections were cut. H&E and immunofluorescences were performed and slides were scanned using NanoZoomer S60 Digital slide scanner and NDP.view2 software (Hamamatsu).

### Immunofluorescence

Tissue sections were submitted to hydration in xylene and a graded alcohol series, slides were steamed in 10 mM citrate buffer (pH 6) for 40 minutes for antigen retrieval. Fixed 3D cultures were permeabilized using 0.5% Triton X-100 in PBS for 20 minutes. Both sections and co-cultures were blocked with 3% Bovine Serum Albumin (BSA, Fisher Bioreagents) and 5% normal goat serum (NGS, Gibco PCN5000) in PBS for 1 hour at room temperature. Primary antibodies (**Suppl Table 2**) were incubated overnight at 4°C, followed by washing and secondary antibodies (Invitrogen, 1:1000) incubation at room temperature for 1 hour in the dark. Slides were mounted with ProLong Gold Antifade reagent with DAPI (Invitrogen, P36931) and images were obtained using Leica Software on a Leica SPE confocal microscope. All quantifications were done using double blind method (quantification of coded samples and de-codification upon completion to interpret the results across multiple animals).

### Whole-mount staining

Calvaria bones were fixed overnight at 4°C in 4% PFA, blocked overnight with agitation in 3% BSA-5% NGS, followed by a 24 hour incubation at 4°C with agitation with primary antibodies (**Suppl Table 2**), 10 hours of washing, secondary antibodies (Invitrogen, 1:1000) overnight incubation at 4°C with agitation and 10 hours of washing. Images were obtained using Leica Software on a Leica SPE confocal microscope.

Sternal bones were collected and transected with a surgical blade into 2–3 fragments, 10 minutes after retro-orbital injection of CD144/VE-cadherin, CD31 and Sca-1 antibodies (1:20). The fragments were bisected sagittally for the bone marrow cavity to be exposed, fixed in 4% PFA and immunofluorescence staining for CK8/18 was performed as above described. Images were acquired using a ZEISS AXIO examiner D1 microscope (Zeiss) with a confocal scanner unit, CSUX1CU (Yokogawa).

### Quantitative PCR

Sorted cells were processed using Cell-to-CT 1-Step Power SYBR Green kit (Invitrogen, A25600) and primers from **Supplementary Table 1**. GAPDH was used as housekeeping control for all experiments.

### Human samples

The BM plasma samples analyzed for TGFb2 and BMP7 were selected from early breast cancer patients included in the SATT study^10^. In this study the patients were monitored for DTCs 2-3 months (BM1) and 8-9 months (BM2) after completion of adjuvant anthracycline-containing chemotherapy. If DTC positive at 8-9 months, the patients received secondary chemotherapy intervention with docetaxel, followed by subsequent DTC monitoring 1 (BM3) and 12 months (BM4) after docetaxel. Patients with disappearance of DTCs (80%) experienced excellent prognosis compared to the patients with DTC persistence (20%)^10^. The BM aspirates (from posterior iliac crest bilaterally) collected for DTC analyses were diluted 1:1 in PBS and separated by density centrifugation using Lymphoprep (Axis-Shield, Oslo, Norway) as previously described^10^. Later on in the inclusion period the plasma supernatant after density centrifugation was collected (if possible) and stored initially at −20 °C, and later transferred to −80 °C. From the available BM plasma samples, we initially performed a limited analysis of plasma samples from patients being DTC positive at either BM1 or BM2, or experiencing systemic relapse without DTC presence. Of these, 55 had ER positive disease, including 30 patients receiving secondary intervention with docetaxel chemotherapy, and 25 patients with no secondary treatment intervention.

### ELISA

Mouse BM supernatant was collected after BM flush centrifugation and stored at −80°C with phosphatase and protease inhibitors (ThermoScientific). TGFb and pro-inflammatory ELISAs were performed by Eve Technologies Corp. (Calgary, Alberta) using the Luminex™ 100 and 200 systems (Luminex, Austin, TX, USA), Eve Technologies’ TFG-β 3-Plex Discovery Assay® (MilliporeSigma, Burlington, Massachusetts, USA) and Eve Technologies’ Mouse Focused 10-Plex Discovery Assay® (MilliporeSigma, Burlington, Massachusetts, USA). BMP7 ELISA was performed using RayBio kit (RayBio, ELM-BMP7) following manufactures instructions.

Patient BM supernatants were analyzed for TGFb2 and BMP7 by multiplex ELISA at Human Immune Monitoring Center (HIMC) at Mount Sinai. ELISA assays were performed using human TGFb2 (R & D System, DY302) and BMP7 (R & D System, DY354) kits following manufacture’s recommendation. Data were analyzed using a Four Parameter Logistic Fit (4PL) method.

### Cell culture

E0771 (CH3 BioSystems), E0771-GFP (gift from John Condeelis’ lab) and MMTV-PyMT-CFP (gift from Jay Debnath) breast cancer cell lines were cultured in RPMI (Gibco) supplemented with 10% foetal bovine serum (Gemini), 10mM HEPES (Corning), 100 units/ml penicillin and 100ug/ml streptomycin (Corning). The tumorigenic HEp3 (T-HEp3)^57^ head and neck squamous cell carcinoma (HNSCC) PDX line and Ct MSCs and rMSCs^39^ were generated and maintained as described previously.

T-HEp3 cells were engineered to express an ELK-GAL4::hrGFP^40^, p38 shuttle-Clover^41^ and p27K^−^ mVenus^42^ biosensors. T-HEp3 with the biosensors were reversely transfected with siRNAs (siTGFBRIII, Invitrogen 32274204; siBMPRII, Oncogene SR300456B) using Lipofectamine RNAiMAX transfection reagent (Invitrogen) according to the manufacturer’s instructions.

Trans-well co-cultures were performed using permeable trans-well assays (Corning), in which transfected cells were plated in the bottom of the well and sorted Nestin-GFP^−or^ ^+^ cells in the permeable inserts.

### Low-density 3D organoid co-cultures

500-1000 cells (E0771 only or with MSCs) were seeded in 400µl assay medium (each cell line media with reduced FBS content (2-5%) plus 2% matrigel) in 8-well chamber slides (Falcon) on top of 50µl of growth factor-reduced matrigel (Corning). Cultures were treated every 24 hours starting at day 0 with TGFβ2 (R&D), BMP7 (R&D) or BM-conditioned media (BM-CM). Single cells, doublets and clusters were quantified after 4 days and the cultures were fixed with 4% paraformaldehyde (PFA) for 20 minutes.

### Western blot

Samples were collected in RIPA buffer and protein concentrations were calculated using Coomassie Plus protein assay (Thermo Scientific) and a standard BSA curve. Samples were then boiled for 8 minutes at 95°C in sample buffer (0.04 M Tris-HCl pH 6.8, 1% SDS, 1% β – mercaptoethanol and 10% glycerol). 8–10% SDS–PAGE gradient gels were run in running buffer (25 mM Tris, 190 mM glycine, 0.1% SDS) and transferred to PVDF membranes in transfer buffer (25 mM Tris, 190 mM glycine, 20% methanol). Membranes were then blocked in 5% milk in TBS-T (Tris-buffered saline containing Tween-20) buffer. Primary antibodies (**Supplementary Table 2**) were incubated overnight at 4°C. Following washing with TBS-T buffer, HRP-conjugated secondary antibodies were left at room temperature for 1 hour. Western blot development was done using Amersham ECL Western Blot Detection (GE, RPN 2106) and GE ImageQuant LAS 4010.

### Statistical analysis

Sample sizes were chosen empirically and no exclusion criteria were applied. The investigators were not blinded to allocation during experiments but quantifications were done in coded samples to reduce operator bias. Statistical analyses were done using Prism Software and differences were considered significant if p<0.05. Unless otherwise specified, 3 or more independent experiments were performed, all values were included and median, interquartile range and 2-tailed Mann–Whitney U-tests, incidence and Fisher’s exact test or mean and SEM and 2-tailed Student’s t-tests were performed.

**Supplementary Table 1.**
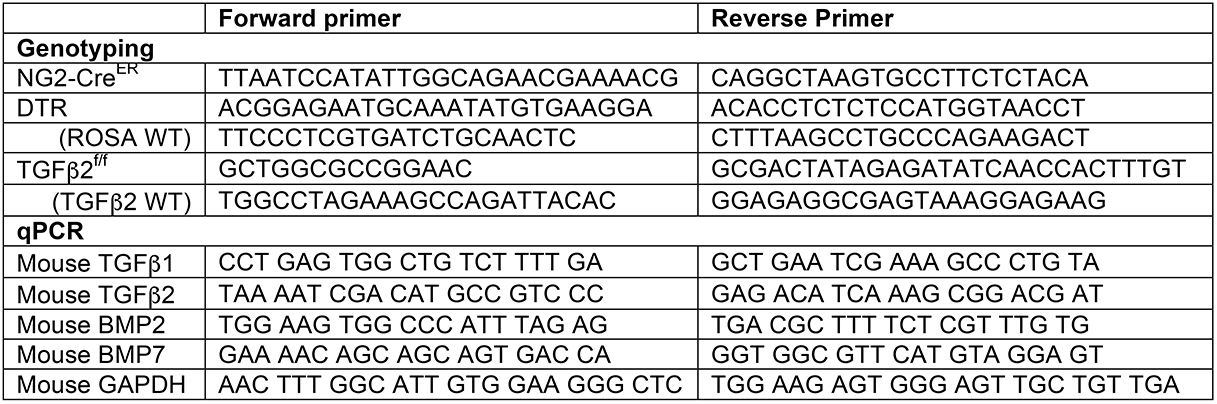
Primers used for genotyping and qPCR.

**Supplementary Table 2.** Antibodies.

|  | Manufacture, Clone/Ref | Conditions |
| --- | --- | --- |
| **FACS** |  |  |
| FITC anti-mouse CD45 | Biolegend, 30-F11 | 1:100, 15min at 4°C in the dark |
| FITC anti-mouse Ter119 | Biolegend, TER-119 | 1:100, 15min at 4°C in the dark |
| FITC anti-mouse CD31 | Biolegend, MEC13.3 | 1:100, 15min at 4°C in the dark |
| APC anti-mouse CD140a (PDGFRa) | Biolegend, APA5 | 1:100, 15min at 4°C in the dark |
| PE anti-mouse CD51 | Biolegend, RMV-7 | 1:100, 15min at 4°C in the dark |
| APC anti-Ki67 | eBiosciences, SolA15 | 1:100, 15min at 4°C in the dark |
| PE anti-p27 | Cell Signaling, D69C12 | 1:100, 15min at 4°C in the dark |
| **IF** |  |  |
| GFP | Aves, 1020 | 1:100, ON (IF) / 24h (whole mount) at 4°C |
| NG2 | R&D, MAB6689 | 1:100, ON (IF) / 24h (whole mount) at 4°C |
| CD31 | Millipore, clone 2H8 | 1:100, ON (IF) / 24h (whole mount) at 4°C |
| p27 | Cell Signaling, D69C12 | 1:100, ON at 4°C |
| Ki67 | Invitrogen, SolA15 | 1:100, ON at 4°C |
| CK8/18 | Progen Biotechnik, GP11 | 1:100, ON at 4°C |
| FITC anti-mouse Sca-1 | Biolegend, D7 | 1:20, 10minutes (retro-orbital) |
| Alexa-647 anti-VE-cadherin | Biolegend, BV13 | 1:20, 10minutes (retro-orbital) |
| Alexa-647 anti-CD31 | Biolegend, MEC13.3 | 1:20, 10minutes (retro-orbital) |
| TGFβ2 | Abcam, ab36495 | 1:100, ON at 4°C |
| BMP7 | Abcam, ab56023 | 1:50, ON at 4°C |
| vE-cadherin | eBioscience, eBioBV13 | 1:100, ON at 4°C |
| cCas3 | Cell Signaling, D175 | 1:100, ON at 4°C |
| pRb | Cell Signaling, D20B12 | 1:100, ON at 4°C |
| pATF2 | Cell Signaling, 9221 | 1:50, ON at 4°C |
| pHistone H3 (Ser10) | Cell Signaling | 1:100, ON at 4°C |
| **WB** |  |  |
| p-SMAD 1/5 | Cell Signaling, 41D10 | 1:500, ON at 4°C |
| p-SMAD 2 | Cell Signaling, 138D4 | 1:500, ON at 4°C |
| pATF2 | Cell Signaling, 9221 | 1:1000, ON at 4°C |
| p27 | Cell Signaling, D69C12 | 1:500, ON at 4°C |
| α-tubulin | Abcam, ab15568 | 1:2000, ON at 4°C |
| HRP horse-anti-mouse IgG | Vector, PI2000 | 1:1000, 1h at RT |
| HRP goat-anti-rabbit IgG | Vector, PI1000 | 1:1000, 1h at RT |

## Acknowledgments

We thank the Aguirre-Ghiso lab for helpful discussions and thank the expertise and assistance of the Dean’s Flow Cytometry CoRE, Microscope CoRE and Small Animal Imaging CoRE at BioMedical Engineering and Imaging Institute, Icahn School of Medicine at Mount Sinai. We are grateful to Ms. Colette Prophete (Frenette’s lab) for mouse husbandry.

This work was supported by The National Institute of Health /National Cancer Institute (CA109182, CA216248, CA218024, CA196521 to J.A.A-G. and HL069438, DK056638, DK116312 and DK112976 to P.S.F) and the Samuel Waxman Cancer Research Foundation Tumor Dormancy Program. A.R.N. was funded by Portuguese Foundation for Science and Technology (SFRH/BD/100380/2014). E.R. was funded by the MD/PhD program of the University of Lyon and the Ecole Normale Supérieure of Lyon. JJBC was funded by the Susan G. Komen Career Catalyst Research (CCR18547848), the NCI Career Transition Award (K22CA196750) and Tisch Cancer Institute NIH Cancer Center Grant (P30-CA196521). AB was supported by a grant from Instituto Serrapilheira/Serra-1708-15285. M.A. was funded by NIH (HL126705, CA218578, R01HL145064) and American Heart Association (17GRNT33650018).

Thank you to Maria Rypdal, Arne V. Pladsen and Ole C. Lingjærde for preparing the BC DTC material and data. The BC DTC work in Oslo has received funding from The Norwegian Cancer Association and Norwegian Health Region South-East (H.G.R. and B.N).

